# A Small Genome Amidst the Giants: Evidence of Genome Reduction in a Small Tubulinid Free-Living Amoeba

**DOI:** 10.1101/2023.12.07.570689

**Authors:** Yonas I. Tekle, Hanna Tefera

## Abstract

This study investigates the genomic characteristics of *Echinamoeba silvestris*, a small-sized amoeba within the Tubulinea clade of the Amoebozoa supergroup. Despite Tubulinea’s significance in various fields, genomic data for this clade have been scarce*. E. silvestris* presents the smallest free-living amoeba genome within Tubulinea and Amoebozoa to date. Comparative analysis reveals intriguing parallels with parasitic lineages in terms of genome size and predicted gene numbers, emphasizing the need to understand the consequences of reduced genomes in free-living amoebae. Functional categorization of predicted genes in *E. silvestris* shows similar percentages of ortholog groups to other amoebae in various categories, but a distinctive feature is the extensive gene contraction in orphan (ORFan) genes and those involved in biological processes. Notably, among the few genes that underwent expansion, none are related to cellular components, suggesting adaptive processes that streamline biological processes and cellular components for efficiency and energy conservation. The investigation delves into genomic structural evidence, including gene content and repetitive elements, illuminating the distinctive genomic traits of *E. silvestris* and providing reinforcement for its compact genome size. Overall, this research underscores the diversity within Tubulinea, highlights knowledge gaps in Amoebozoa genomics, and positions *E. silvestris* as a valuable addition to genomic datasets, prompting further exploration of complexities in Amoebozoa diversity and genome evolution.

## Introduction

The supergroup Amoebozoa is composed of microbial eukaryotes of predominantly amoeboid protists (Cavalier-Smith, et al. 2004; Smirnov, et al. 2011; Tekle, et al. 2008). It encompasses three major lineages including Discosea, Tubulinea and Evosea (Tekle, et al. 2022b). Amongst this the Tubulinea is the most robust clade recovered in most molecular phylogenetic studies and most of its members share morphological recognizable tubular pseudopodia (Lahr, et al. 2013; Smirnov, et al. 2011). Tubulinea includes diverse members including notable members such as the paleontological important testate amoebae that form microfossil and the classic textbook example amoeba genus (e.g., *Amoeba proteus*) whose members are presumed to include the largest genome of all living things (Friz 1968; Porter, et al. 2003; Smirnov, et al. 2011). It also includes some newly added obscure amoebae members such as *Trichosphaerium* considered to have alternation of generation as well as small sized amoeba commonly found in human environments such as hospitals causing health concern to humans (Delafont, et al. 2018; Schaudinn 1899; Sheehan and Banner 1973). Despite the familiarity, the health, ecological and paleontological importance of this clade, genome data representing this clade is scanty.

Genome data in the Amoebozoa is steadily growing although majority of the available genomes are from restricted genera that include model organisms or those implicated in human health (Clarke, et al. 2013; Eichinger, et al. 2005; Loftus, et al. 2005). Most genome data that are publicly available for the Amoebozoa belong to the clade Evosea (24 total of these 12 are annotated) and Discosea (18 total, only two annotated genomes) (Chelkha, et al. 2020; Tekle, et al. 2022a; Tekle, et al. 2021b; Zahonova, et al. 2022). Tubulinea is only represented by two genomes including *Vermamoeba* (Chelkha, et al. 2020) and the recently published *Trichosphaerium* genome (Tekle, et al. 2022a). The genome size of the majority of the published Amoebozoa genomes are quite small (average 43 MB) compared to what is reported for the supergroup. A member of Tubulinea, *Amoeba dubai,* is reported to have the largest genome size of all known living organisms (200x larger than the human genome) (Friz 1968). However, this observation is only based on qualitative data and requires further verification. In this study we present the draft genome of an isolate belonging to a clade comprising small tubulinid amoebae characterized by special adaptation and posing potential health concern to human.

*Echinamoeba* is a genus of flat, diminutive amoebae that lack protective coverings, are single-nucleated, and exhibit distinctive, spine-like subpseudopodia (Baumgartner, et al. 2003; Page 1975; Page 1967; Rohr, et al. 1998). Species belonging to this genus are described from thermal and non-thermal natural environments, as well as from hospital hot water systems. The closest relative of the genus is *Vermamoeba vermiformis*, within the order Echinamoebida, which is known for its inhabitance in human environments and pathogenicity (Page 1975; Page 1967; Rohr, et al. 1998). Members of Echinamoebida are interesting groups of amoebae adapted to extreme environments and their potential impact on human health is a concern (Baumgartner, et al. 2003; Fields, et al. 1989). *V. vermiformis* is the only and the first genome sequenced for the clades Echinamoebida and Tubulinea, respectively (Chelkha, et al. 2020). Comparative genome study of these amoebae will unravel important insights into their evolution, adaptation, and potential pathogenicity in humans. Draft genome analysis of *Echinamoeba silvestris* CCAP 1519/1 reveals that it is one of the smallest genome sizes among free-living amoebae with genome data. The number of predicted gene models in *E. silvestris* is comparable to those parasitic amoebae with reduced genomes (Loftus, et al. 2005). The genomic content and architecture and evolution related to genome size and complexity of this amoebae with other amoebozoans is discussed.

## Results

### Genome architecture and gene prediction of *Echinamoeba silvestris CCAP 1519/1*

Using three sequencing technologies, we generated over 161.4 million sequencing reads (>100× coverage), comprising 151.3 million Illumina short reads and long reads totaling 10.1 (3.26 Gb) and 9.0 (16.68 Gb) million, respectively, from Oxford Nanopore and PacBio. These genomic data were derived from amplified DNA nuclei pellets. To ensure data integrity, we implemented a series of bioinformatics and manual curation steps described in the methods to eliminate contamination from known food bacteria or associated entities, symbionts (e.g., viruses, archaea), and other environmental contaminants. From these decontaminated data, we successfully generated a draft genome of *Echinamoeba silvestris*, totaling 27.1 megabase pairs (MB) (Table 1). The assembly comprises a total of 282 scaffolds, with an average scaffold length of 95,403 base pairs (bps). *E. silvestris*’s genome falls within the higher GC content range (47.77%) among amoebozoan genomes (Table 1).

**Table 1.** Genomic composition and gene model repertoire of *Echinamoeba silvestris* CCAP 1519/1.

| Feature | Composition |
| --- | --- |
| Genome size (bp) | 27.1MB |
| GC content (%) | 47.77 |
| DNA scaffolds | 284 |
| Longest scaffold length (bp) | 3,419,598 |
| Shortest scaffold length (bp) | 1,192 |
| Mean scaffold length (bp) | 95,403 |
| N50 (bp) / L50 | 984,026 |
| Total number of predicted transcripts | 8,327 |
| Proportion of transcripts with a size $\geq 300$ bp | 6,034 |
| Genes assigned to Cluster Orthologous Groups (COGs) | 8,327 |
| Non-ORFan genes | 7,017 |
| ORFan genes | 1,310 |
| Mean number of introns/gene | 0.9 |
| Mean number of exons/gene | 1.9 |
| Mean intron size (bp) | 164.7 |
| Mean exon size (bp) | 842.1 |
| Gene model BUSCO completeness (complete + partial) | 83.2% |

The overall genomic content of *E. silvestris* including introns, exon and gene numbers are smaller compared to published free-living amoebozoan genomes (Clarke, et al. 2013; Eichinger, et al. 2005; Loftus, et al. 2005). The prediction of gene models of *E. silvestris* was aided by transcriptome data from a closest relative *E. exundans* and a published genome of an amoeba, *Acanthamoeba castellanii*. Using this approach, we generated a total of 8,327 gene models. Almost all transcripts obtained from the *E. exundans* transcriptome were found in the draft genome of *E. silvestris* with high similarity percentage matches. In addition to this the majority, 84.3% (7,017), of gene models were assigned to well-known biological processes in the Clusters of Orthologous Groups of proteins (COGs) database. These includes the Cellular Processes and Signaling (27.8%), Metabolism (21.3%), Information, Storage and Processing (17.9%), categories (Table S1). The remaining 20.9% COG category included genes that are poorly characterized or of unknown function (see Table S1). Despite its small genome, *E. silvestris* possesses a similar number of common genes, including ribosomal and cytoskeletal genes (data not shown), as well as genes involved in meiosis (see below), comparable to other amoebae.

The *E. silvestris* draft genome comprises a very small fraction of ORFans (1.86%, Table 1). ORFans are genes that lack BLAST hits in the NCBI GenBank database and likely represent novel or unique genes specific to the amoeba. While the exact functions of ORFans remain unknown, some of these genes are expressed, as evidenced by their presence in the *E. exundans* transcriptome data. A preliminary functional exploration of some of these genes, using InterPro, has unveiled some common protein domains in some of these ORFans. These domains include those involved in protein–protein interactions (e.g., Lucine-rich and Ankyrin-repeats) and membrane-associated proteins (e.g., GRAM domain).

### Taxonomic distribution and interdomain LGT of predicted gene models in *E. silvestris*

Similar to other amoebae genomes, the *E. silvestris* genome displays a diverse distribution of gene models across various taxonomic groups. The majority (70.13%) of the gene models matched eukaryotic genes (Fig. 1). Among other living domains, the largest proportion (26.6% - 2,215 genes) show highest similarities to bacteria, while only a small fraction shows similarities to archaeal genes (0.8%, 67 genes) (Fig. 1). An even smaller percentage of gene models (0.6%, 49 genes) show similarity to viral genes. This latter set, non-eukaryote matching genes, makes up core components of cellular (signaling and metabolism) and information storage and processing (Fig. S1). Expression of some of these gene models with high similarity to bacterial, archaeal and viruses have been detected in transcriptome data of *E. exundans* (Table S2).

**Fig. 1.**
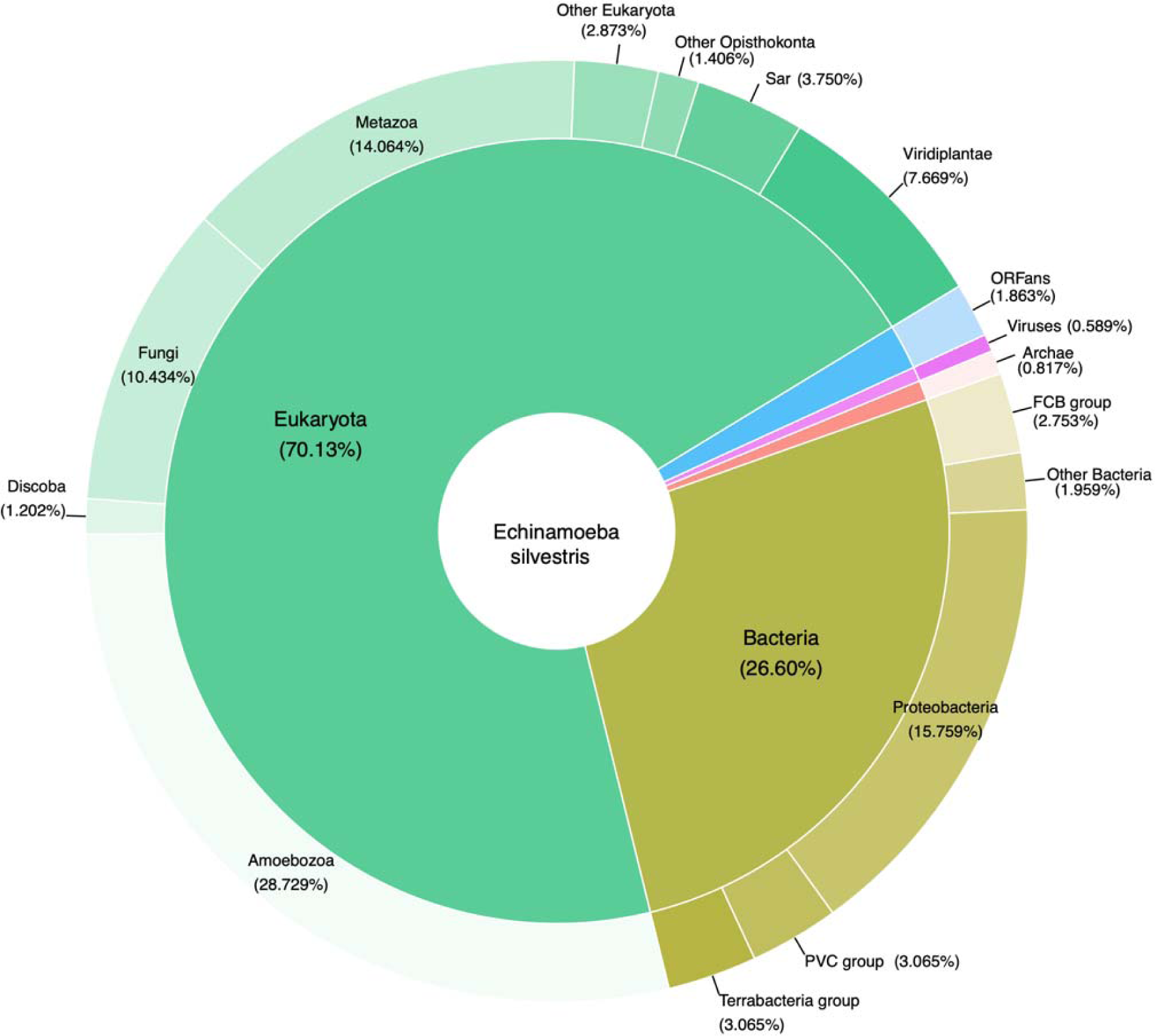
Taxonomic classification of predicted proteins deduced from *Echinamoeba silvestris* draft genome.

To assess evidence for lateral gene transfer (LGT) of the non-eukaryotic predicted gene model we conducted alien index analyses and phylogenetic analysis on selected genes (Fig. S2). Based on BLAST similarities and alien index analyses, we identified a large number of LGT candidates from bacteria and archaea. The likely donors of the putative LGTs in bacteria come from diverse taxonomic groups but most prominently (66%) from Pseudomonadota (Proteobacteria). Other major potential donors include PVC group (10.7%), FCB group (7.2%) and Terrabacteria group (4.7%) (see Fig. S2*a,b*; Table S2).

Among the 67-archaeal origin genes found in the *E. silvestris* draft genome, only 2 genes had an alien index above the threshold (Table S2). A phylogenetic analysis of the putative archaeal LGT showed that both *E. silvestris* and *Vermamoeba vermiformis* acquired a similar gene that encodes for lactate dehydrogenase-like protein (Fig. S2*c*). The likely donor of these putative gene is Euryarchaeota.

Similarly, *E. silvestris* have a dozen of putative LGTs, with alien indexes above the threshold, of viral origin (Fig. S2*d*, Table S2). Majority of these putative LGTs have their origins from giant virus clade Mimiviridae, which include *Acanthamoeba polyphaga* mimivirus, Klosneuvirus KNV1, *Hyperionvirus* sp. and Fadolivirus. The others unclassified giant viruses including Kaumoebavirus and Pithovirus (Table S2). The former giant virus has been reported to be associated with a closest relative of *V. vermiformis* (Bajrai, et al. 2016).

### Comparative genomics and genes involved in sexual life cycle

Comparative genome analysis of amoebae representing the three clades of amoebae, Tubulinea (*E. silvestris, V. vermiformis* and *Trichosphaerium* sp.), Discosea (*A. castellanii* and *C. minus*) and Evosea (*E. histolytica* and *D. discoideum*) reveal interesting results (table 2). *E. silvestris* is the smallest free-living genome to the exclusion of the parasitic *E. histolytica*. More interestingly *E. silvestris* has significantly lower number of predicted gene models comparable to the parasitic amoebae (table 2). Despite variations in morphological size, most sequenced free-living amoebae genomes consistently exhibit an average of ∼17,500 genes (ranging from 13,315 to 27,369). The analysis of genome size versus the number of predicted gene models reveals robust correlations (Fig. 2).

**Table 2.** Genomic comparison of diverse Amoebozoan genomes.

| Species | Genome Size - MB | (%) GC content | Predicted genes (CDs) | ORFan genes | Repeat Elements | Mean no. introns/gene | Mean no. exons/gene |
| --- | --- | --- | --- | --- | --- | --- | --- |
| *E. silvestris* | 27.1 | 47.77 | 8,327 | 155 | 6.83 | 0.9 | 1.9 |
| *V. vermiformis* | 59.55 | 41.7 | 22,473 | 7,220 | 9.62 | 2.7 | 3.6 |
| *Trichosphaerium* sp. | 70.8 | 37.7 | 27,369 | 7,489 | 14.54 | 8.7 | 7.7 |
| *E. histolytica* | 20.15 | 24.3 | 8163 | 2 | 30.78 | NA | 0.77 |
| *D. discoideum* | 34.21 | 22.5 | 13315 | 63 | 34.34 | NA | 0.46 |
| *C. minus* | 50.5 | 26.33 | 19,925 | 5,723 | 12.84 | 4.7 | 5.7 |
| *A. castellanii* | 42 | 57.9 | 14,969 | 0 | 4.04 | 6.2 | 7.1 |

**Fig. 2.**
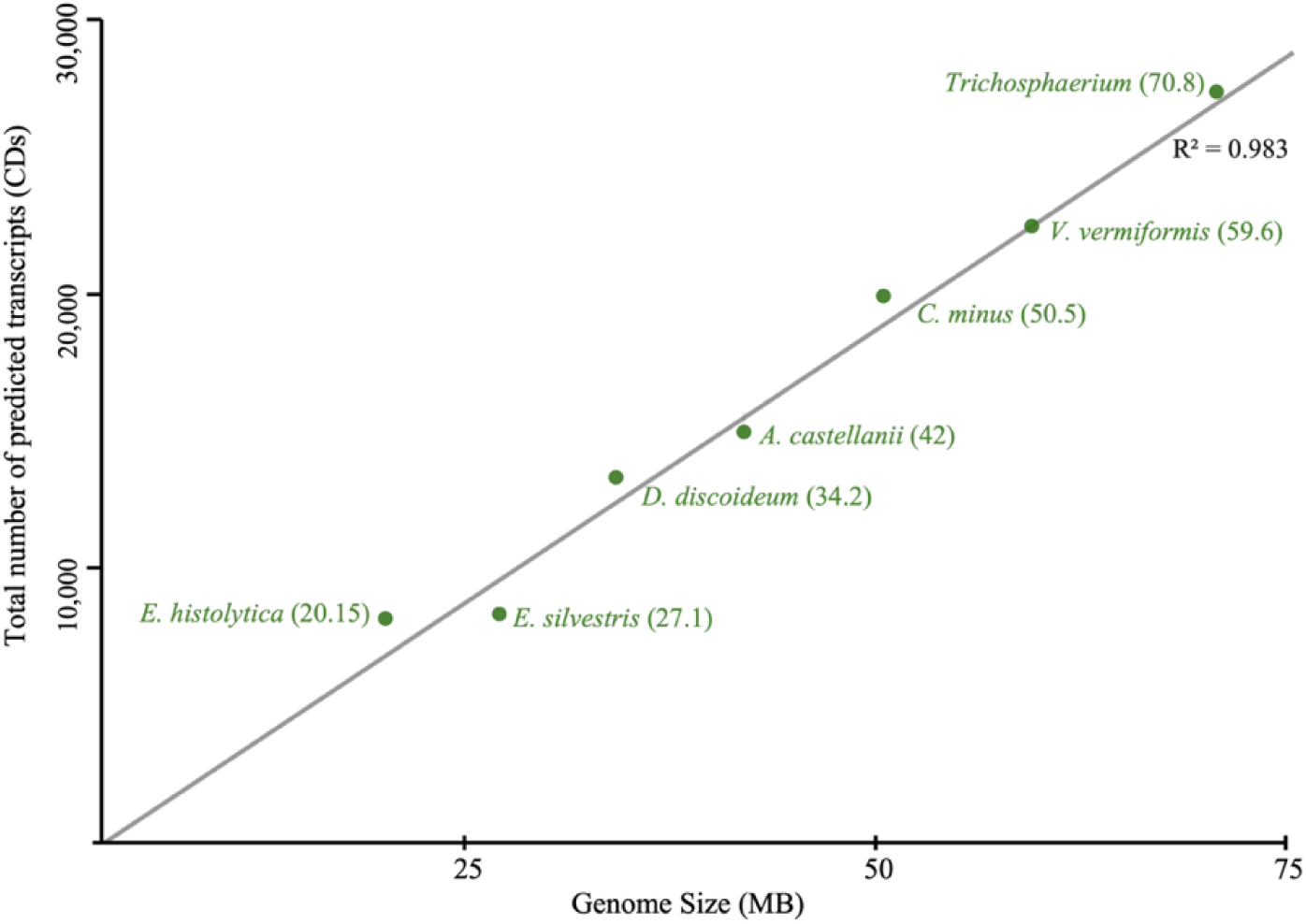
Correlation between genome size and number of predicted genes in Amoebozoans.

A detailed examination of the genome architecture reveals that *E. silvestris* has the lowest average number of introns per gene compared to its closest relatives in the Tubulinea clade and other members of Discosea (table 2). Intron information for members of Evosea is not available. Analyzing the number of ORFan genes among Tubulinea members, *E. silvestris* emerges with the lowest total percentage. However, a cross-amoebae comparison of ORFan genes proves challenging due to the comprehensive classification of genes in model and parasitic organisms resulting from extensive studies. Nevertheless, it is noteworthy that among Tubulinea members whose genomes are assembled and annotated with a similar approach, *E. silvestris* is shown to have a significantly lower number of ORFan genes (table 2). Similarly, *E. silvestris* exhibits the lowest percentage of repetitive elements in its genome (table 2).

The analyzed amoebae display a range of GC content in their genomes. In general, members of Tubulinea exhibit higher average GC content, while members of Discosea and Evosea have lower GC content, with one exception: *A. castellanii*, which has the highest GC content among all studied amoebozoans so far (table 2).

Comparative genomics also reveals that *E. silvestris* has experienced extensive gene family contraction in comparison to other amoebae (Fig. S3 and Table S2). Similarly, the number of gene families undergoing expansion is notably lower in *E. silvestris* (Fig. S3). The majority of gene families undergoing expansion and contraction are associated with biological processes and molecular functions, with only a few classified under cellular components (see Table S3). Among the expanded gene families are those involved in nuclear import, cell movement and division, as well as proteins contributing to apoptosis and pathogen defense (Table S3).

To explore the sexuality of *E. silvestris* and assess genome completeness using well-known genes in amoebae genomes, we conducted a gene inventory analysis of meiosis-specific genes (Table S4). *E. silvestris* possesses almost all known meiosis-specific genes in amoebae and other eukaryotes (Table S4). Two genes, *MND1* and *HOP2*, were not found in this draft genome. Although the absence of these genes might be due to the incompleteness of the draft genome, it is common to observe the absence of one or a few meiosis genes in several amoebae genomes, including notable cases such as the absence of *SPO11* and *DMC1* in *D. discoideum* (Table S4).

## Discussion

### *E. silvestris* the smallest free living amoebae genome to date

Comparative genomics of Amoebozoans, encompassing understudied groups, is yielding valuable insights into the evolution and genetic diversity within this group (Chelkha, et al. 2020; Tekle, et al. 2021a; Tekle, et al. 2022a; Tekle, et al. 2021b; Tekle, et al. 2022b; Zahonova, et al. 2022). Our comprehension of Amoebozoa supergroup genomes has hitherto been constrained by well-studied model organisms and other traditional (qualitative) genome data resources, often lacking in-depth details (Eichinger, et al. 2005; Glockner and Noegel 2013; Loftus, et al. 2005). Here, we present a draft genome of an amoeba belonging to a group comprised of small-sized amoebae, encompassing lineages with potential pathogenicity and adaptation to extreme temperatures (Baumgartner, et al. 2003; Delafont, et al. 2018). This study contributes a third genome dataset to the Tubulinea clade, the least examined clade in genomic studies of the Amoebozoa supergroup (Chelkha, et al. 2020; Tekle, et al. 2022a). Tubulinea encompasses diverse lineages of amoebae, exhibiting variations in behavior, morphology, and ecology (Smirnov, et al. 2011). Traditionally recognized as a clade possessing large genome sized members (Friz 1968), our study reveals the smallest free-living amoeba genome within the Tubulinea clade. This underscores the substantial diversity within the Tubulinea clade and emphasizes the existing knowledge gaps in Amoebozoa genomics and diversity.

While the available data on the nature and diversity of genomes in Tubulinea is limited, our discovery of the small genome observed in *E. silvestris*, comparable to a parasitic lineage, is intriguing in its own right. Among the analyzed amoebae genomes, the only genomes closest in size and number of predicted gene models are those of the parasitic genus *Entamoeba* (table 2, see below). The average genome size in free-living amoebae, regardless of morphology or life history complexity, is 43 MB (see table 2). The genome of *E. silvestris* is approximately half the size of its closest relative, *V. vermiformis*, and about 20% smaller compared to the smallest (before the current study) free-living amoeba, *D. discoideum* (table 2, Fig. 2). The consequences of reduced or small genomes and their evolutionary significance in free-living amoebae and other microbial eukaryotes are not well understood due to limited data (Zhang, et al. 2022).

A previous report showed that cell size is correlated with genome size across many taxonomic groups (Bennett and Leitch 2005; Gregory 2005). *E. silvestris* (∼11µm) is the smallest in size compared to the amoebae examined in this study. This observation supports these previous reports; however, cell sizes in the studied amoebae have a close range (ranging 10-60 µm), with some of them only a few microns apart and still displaying large proportions of genomic size variations among cells of similar size. Therefore, more data is needed to examine the correlation between cell and genome size in amoebae. It is worth noting that the division (growth) rate has been observed to correlate with genome size, and the trend suggests that organisms with smaller genomes tend to exhibit higher division rates compared to those with larger genomes (Bennett and Leitch 2005; Gregory 2005). Despite possessing the smallest genome, *E. silvestris* stands out as one of the slowest-dividing amoebae among the free-living amoebae we cultivate in our laboratories under similar conditions. This adds complexity to the association between genome size and cellular characteristics. The association between genome size and organismal complexity is one of the contentions and complex subjects in eukaryotic evolution. This is generally called the C-value enigma and is mostly based on mostly observations obtained in macrobial (animals and plants) organisms (Thomas 1971). With more genome data from microbial eukaryotes, it is possible to gain insights into this intriguing topic, considering factors such as body size, metabolism, developmental rate, and geographical distribution.

### Genome architectural evidence for small genome size in *E. silvestris*

The intricacies of genome evolution, particularly concerning size, in eukaryotes are multifaceted. Eukaryotic genomes exhibit a substantial size range compared to prokaryotes, a phenomenon influenced by various additional genomic features and evolutionary forces shaping their development. Determinants of genome size in eukaryotes include factors such as functional complexity (gene content), repetitive sequences, transposable elements, ploidy level, non-coding DNA, and introns (Elliott and Gregory 2015). Additionally, the evolutionary history, marked by events such as duplications, deletions, rearrangements, along with environmental factors, serves as additional determinants influencing the evolution of genome size in eukaryotes. In this study, we conducted a comparative genomics analysis on the draft genome of *E. silvestris* to elucidate specific aspects related to the evolution of genome size in amoebae.

Our findings provide multiple lines of evidence supporting the observed small genome size in *E. silvestris*. Gene content emerges as a prominent contributor to genome size in amoebozoans [table 2, (Delafont, et al. 2018; Eichinger, et al. 2005; Tekle, et al. 2022a; Tekle, et al. 2021b)] and other microbial eukaryotes (Elliott and Gregory 2015). The total predicted number of genes in amoebae exhibits a robust correlation with genome size (Fig. 2). Notably, *E. silvestris* has the lowest number of predicted genes among the free-living amoebae examined in our study (table 2). While limited publicly available genome data exist for amoebae, our preliminary data on other diverse amoebae with comparable size appear to have substantially more predicted genes than *E. silvestris* (Tekle, unpubl. data).

Surprisingly, the number of predicted genes in *E. silvestris* is comparable to that of parasitic amoebae (table 2). This observation is intriguing considering that parasitic lineages are generally considered to have reduced genomes due to adaptive processes associated with optimizing the parasitic lifestyle (Corradi and Slamovits 2011; Jackson 2015; Keeling and Slamovits 2005). Such processes include evolutionary pressures favoring the loss of non-essential genes and pathways, driven by factors such as host dependency, energy efficiency, symbiosis, specialization, and reliance on host functions. Parasitic lineages of amoebae, exemplified by *Entamoeba histolytica*, also lack mitochondria, further impacting their overall functional genetic composition.

Functional categorization of predicted genes in *E. silvestris* reveals similar percentages of genes assigned to cellular processes and signaling, metabolism, information storage and processing, as observed in other amoebae (Table S1). *E. silvestris* also have similar number of essential genes such as cytoskeleton and ribosomal as well as those genes involved in meiosis (Table S4). However, *E. silvestris* appears to have undergone extensive gene contraction relative to other amoebae (Fig. S3, Table S3). Two noteworthy observations include the substantial number of gene contractions in genes involved in biological processes, and among the few genes that underwent expansion, none are related to cellular components (Table S3). It is plausible that the diminutive size of the amoeba has led to adaptive processes streamlining biological processes and cellular components for efficiency and energy conservation.

In our investigation of gene content within less-explored amoebae genomes, a noteworthy observation emerges from the substantial presence of genes lacking significant homology in public databases, commonly known as ORFans. These ORFans constitute a considerable proportion of amoebae genomes and likely contribute to shaping unique morphological features and other adaptations specific to amoebae (Tekle, et al. 2021b). Notably, *E. silvestris* has a significantly lower percentage of ORFans (4.7%) compared to its closely related species (*V. vermiformis*, 32%, and *Trichosphaerium* sp., 27%) and a distantly related species, *C. minus* (29%), potentially accounting for its smaller genome size. However, it is crucial to interpret these findings with caution, considering that extensively studied and well-characterized genomes, such as those of *E*. *histolytica*, *D. discoideum*, and *A. castellanii*, exhibit minimal numbers of ORFans. Consequently, while our comparative analysis suggests a significant reduction of ORFans in the *E. silvestris* genome compared to its close relatives, whose genome data is collected using a similar approach, further validation is essential as our comprehension deepens with improved amoebae genome characterization. The functional characterization of ORFans, representing a substantial portion of the genome, holds the promise of unraveling intriguing insights into the evolution and diversification of this group.

Our comparative analysis of non-coding and repetitive elements provides additional insights into the genome size evolution of amoebae. Non-coding DNA constitutes a significant contributor to genome size evolution in eukaryotes, encompassing various elements such as promoters, enhancers, silencers, non-coding RNA molecules, intergenic regions, introns, and repetitive elements. While the nature and contribution of non-coding DNA in multicellular eukaryotes are well-documented (Elliott and Gregory 2015), variations may exist across species or taxonomic groups. On average, non-coding DNA constitutes 27% and 51% of the genome in animals and plants, respectively (Elliott and Gregory 2015). In some cases, such as in mammals and vertebrates, the majority of their genome (>90%) is composed of these elements (de Koning, et al. 2011; Mattick and Makunin 2006).The nature and evolution of non-coding regions in genomes of microbial eukaryotes in general, and amoebae in particular, are poorly understood (Elliott and Gregory 2015). Limited studies on repetitive elements, focusing on the parasitic lineages of amoebae, exist, reporting varying types of repetitive elements (Bakre, et al. 2005; Lorenzi, et al. 2008). In this study, we compiled a large database of repetitive elements and conducted a comprehensive comparison within amoebae representing the three major groups. We found consistent results with various types of non-coding elements in amoebae. A detailed report of this study will be published elsewhere. Here, we report the contribution of repetitive elements to genome size in relation to *E. silvestris*. Among the species analyzed, *E. silvestris* has the lowest percentage of repetitive elements in its genome. Members of Evosea (*E. histolytica* and *D. discoideum*) have the highest percentages of repetitive elements (table 2). While a more thorough study is required to investigate the variation observed between the major clades, this result clearly demonstrates that repetitive elements significantly contribute to the genome size evolution of amoebas, similar to multicellular organisms (Elliott and Gregory 2015). The lower percentage of repetitive elements in *E. silvestris* corroborates their contribution to genome size. Repetitive elements are found to positively correlate with genome size in other eukaryotes (Elliott and Gregory 2015), although such a conclusion cannot be reached in amoebae due to limited sampling. Similarly, *E. silvestris* has the lowest mean number of introns per gene, further supporting the observed reduced genome in this amoeba. Similar pattern of genome reduction in repeat elements and other non-coding regions are also known in one of the smallest free-living eukaryotes, green alga, *Ostreococcus tauri* (Derelle, et al. 2006).

### Concluding remarks

In conclusion, our study presents the draft genome of *E. silvestris*, the smallest free-living amoeba genome identified to date. This research contributes to the comparative genomics of amoebozoans, shedding light on the evolution and genetic diversity within this group. The exploration of the Tubulinea clade, one of the least sampled clades in amoebozoan genomics, underscores the significant diversity within this lineage. Despite the limited data on Tubulinea genomes, our findings reveal the unique nature of *E. silvestris*, exhibiting a genome size comparable to parasitic lineages, challenging conventional expectations of genome reduction in parasites. Additionally, our investigation into non-coding and repetitive elements provides insights into the evolution of genome size in amoebae, with *E. silvestris* standing out for its low percentage of repetitive elements. However, the observed correlation between genome size and cellular characteristics, such as division rate, adds complexity to the understanding of the evolutionary significance of genome size in free-living amoebae. Further research is warranted to explore the implications of these findings on amoebae genomics and the broader context of eukaryotic genome evolution.

## Materials and methods

### Genomic DNA collection of Echinamoeba silvestris CCAP 1519/1

Culture of *Echinamoeba silvestris* was obtained from Culture Collection of Algae and Protozoa (CCAP). Subsequent monoculture of this isolate was grown for genomic DNA isolation in plastic petri dishes with bottled spring water (Deer Park®; Nestlé Corp. Glendale, CA) at room temperature supplemented with autoclaved grains of rice. The identification of the isolate was confirmed using morphological and genetic analysis against published literature for the isolate. *E. silvestris* on average is 11 µm in size with locomotive morphology of triangular or elongate forms. It displays the characteristic spine-like subpseudopodia of the genus (Baumgartner, et al. 2003; Page 1975).

Genomic DNA from *E. silvestris* was extracted from monoclonal cultures following a previously published protocol for nuclear extraction and genome amplification (Tekle, et al. 2022a; Tekle, et al. 2021b). The nuclear extraction process began by lysing cleaned adherent cells to release the nuclei. This was achieved by incubating the cells in 6 ml of lysis buffer, composed of sodium phosphate buffer at pH 7.4, 5 mM MgCl2, and 0.1% Triton-X 100, for 2 hours. Subsequently, the lysate containing free nuclei was separated by centrifugation at 500 rpm for 10 minutes at room temperature. The resulting nuclei pellet was resuspended in 0.5 ml of lysis buffer and carefully layered on top of a 12 ml sucrose cushion consisting of 30% sucrose, sodium phosphate buffer at pH 7.4, and 0.5% Triton-X 100. The sucrose cushion played a crucial role in separating the nuclei by capturing small particles (e.g., bacteria) or lightweight lysates (cell parts) during centrifugation. The mixture of lysate and sucrose cushion was then centrifuged at 3200 rpm for 20 minutes at room temperature. The pellet obtained from this step was re-suspended in 1 ml of lysis buffer and subjected to further centrifugation at 10,000 rpm for 1 minute at room temperature. The purified nuclei pellets were carefully collected after removing the supernatant. Whole genome amplification (WGA) nuclei pellets were carried out using the REPLI-g Advanced DNA single-cell kit (QIAGEN; Cat No./ID: 150363), following the manufacturer’s protocol. Finally, the amplified DNA was quantified using the Qubit assay with the dsDNA broad-range kit (Life technologies, Carlsbad, CA, USA).

### Genomic DNA library preparation and sequencing

We applied three different sequencing strategies: Illumina short reads, 10x genomics linked reads, and Oxford Nanopore long read sequencing. For Illumina and Pacbio sequencing, we sent a nucleus pellet amplified gDNA sample to GENEWIZ (Azenta Life Sciences, South Plainfield, NJ) for library preparation and sequencing as described in (Tekle, et al. 2021b). Oxford Nanopore technology (ONT) (Oxford Nanopore Technologies Ltd., United Kingdom) was used to generate long-read sequences in our lab with MinION device using the SQK-RAD004 kit. We constructed the library from 400 µg of amplified nuclear or single cells gDNA and this library mix was added to the flow cell using the SpotON port.

### *De novo* Genome assembly, Contaminant removal

We employed a comprehensive suite of bioinformatics pipelines, building upon our prior work, to assemble and annotate the genome while eliminating contamination (Tekle, et al. 2022a; Tekle, et al. 2021b). Initially, we assembled Nanopore and PacBio long-read sequences, using the Canu assembler. Subsequently, this assembly underwent careful refinement over three iterations, integrating high-quality Illumina reads using Pilon v1.2. To further enhance the assembly quality, we applied Redundans to remove heterozygous genome segments and eliminate short (<1000 bp) contigs. This refined assembly served as the basis for contamination assessment (see below). By aligning trimmed Illumina short reads and Canu-corrected long reads to the polished assembly using minimap2, we determined per-contig (scaffold) coverage. Additionally, we used a diamond-blastx search against the NCBI nt database to assign taxonomic annotations to each scaffold. Scaffolds meeting specific criteria, such as non-eukaryotic taxonomic hits and coverage < 10.0, were flagged as contaminants and excluded from subsequent analyses. The remaining scaffolds were employed for gene prediction (see below). To further scrutinize contamination, we performed a BLASTP search against the NCBI non-redundant protein database (nr) for predicted gene models. Scaffolds with significant BLASTP hits to potential contaminants, particularly those specially linked to bacteria, archaea, viruses, or non-amoeboid eukaryotes, were carefully examined, and removed. The decontaminated scaffolds were analyzed using BUSCO and transcriptome data to confirm that amoeba (eukaryote) genes were preserved.

### Gene prediction, LGT analysis and identification of sex-related genes

We followed the methodology for gene annotation described in (Tekle, et al. 2021b) using transcriptome data from a closely related amoebae *Echinamoeba exundans* and protein sequences from a published genome of a related amoeba, *Acanthamoeba castellanii* (Clarke, et al. 2013). Following the prediction of the finalized gene set, we classified the predicted gene models into Clusters of Orthologous Groups (COGs) categories using EggNOG-mapper, as implemented in OmicsBox v.2.0.29. Functional annotations of coding-gene were obtained from the best BLAST hit with BLASTP (e-value < 1e-3) against the NCBI nr database. Genes that had no hits to nr database were classified as ORFans. Genome annotation quality was evaluated by BUSCO using the Eukaryote odb10 database.

To detect potential lateral gene transfer (LGT) from the draft genome of *E. silvestris,* we checked extracted genes that matched to bacteria, virus, and archaea based on their non-redundant annotations as LGT candidate for further analyses. We performed BLASTP to search these genes against archaea, bacteria and virus genomes that are commonly seen in previous blast results. We then performed an Alien Index (AI) analysis based on the BLASTP results similar to our published methods (Tekle, et al. 2021b) to calculate the AI scores. Genes with AI ≥ 45 were considered as putative LTGs. To further corroborate these results, we built phylogenetic trees for selected putative LTGs in IQ-Tree using the automatic model selection option and 1000 ultrafast bootstrap replicates.

To identify gene models of certain genes important in adaptive behavior (thermal or pathogenicity) and those implicated in sexual reproduction within the draft genome of *E. silvestris*, we performed BLASTP (e-value 1e-15) search of respective genes as in (Wood, et al. 2017). The selected genes were further analyzed using phylogenetic analysis in IQ-Tree as described above to further assess their homology in phylogenetic framework.

### Repetitive elements, gene family and ontogeny analyses

We constructed a comprehensive non-redundant repeat elements database encompassing all Amoebozoans, as outlined in references (Berriman, et al. 2018; Šatović, et al. 2020). To compile this database, repetitive elements identified by three distinct software packages (LTRharvest, TransposonPSI, RepeatModeler) were combined into a unified library for each species. The outcomes of these analyses were categorized using the Repeat classifier script within the RepeatModeler package (Flynn, et al. 2020). These classifications were further consolidated into a master database, eliminating duplicate repeats through VSEARCH (Rognes, et al. 2016). This master database served as the foundation for conducting comparative analyses of repeat elements present in Amoebozoa genomes. The final percentage of repetitive elements within each genome was determined by applying RepeatMasker against this non-redundant database.

For general whole-genome comparative analysis of protein-coding genes, including clustering based on orthology and function, we employed OrthoVenn3 (Wang, et al. 2015), a web-based tool. Additionally, OrthoVenn3 functionalities were utilized for gene family expansion-contraction history analysis, employing the CAFE5 (Mendes, et al. 2021) software.

## Acknowledgments

We would like thank Hann Tran for technical assistance during data collection and analysis.

## Author Contributions

YIT conceived the project, led writing manuscript and helped design experiments and analyses. HT collected data, conducted analyses, and contributed to writing and editing of the manuscript. Both authors have read and approved the manuscript.

## Supporting information legends

**Fig. S1**. Functional categorization of putative HGTs (LGTs) found in *Echinamoeba silvestris* draft genome based on the Cluster Orthologous Groups (COGs) database.

**Fig. S2**. Phylogenetic reconstructions demonstrating putative lateral gene transfers (LGTs) in genome among bacteria (A, B), archaea (C), and giant viruses (D) from *Echinamoeba silvestris*. Clade supports at nodes are ML IQ-TREE 1000 ultrafast bootstrap values. All branches are drawn to scale.

**Fig. S3**. Gene family expansion and contraction based on whole genome coding proteins of amoebae constructed using inbuild function of OrthoVenn3 (Wang, et al. 2015). Pie charts at internodes (shared) and terminal taxon (unique) show number of genes that underwent contraction (negative values - blue) and expansion (positive values - purple).

**Table S1**. Distribution of the gene models of *Echinamoeba silvestris* draft genome in categories of clusters of orthologous groups of proteins (COGs).

**Table S2**. Putative LGT-derived genes in *Echinamoeba silvestris* draft genome with Alien Index above threshold (>45) scores.

**Table S3**. Functional classification of gene families that underwent contraction (sheet1) and expansion (sheet2).

**Table S4**. Meiosis genes inventory in members of Amoebozoa.

